# A unified molecular theory of sweet taste: revisit, update and beyond

**DOI:** 10.1101/2023.08.01.551443

**Authors:** Bo Liu

## Abstract

The molecular mechanism for the generation of sweet taste is still elusive, mainly because there has no common feature revealed imparting sweetness to various sweeteners^1-2^, although many principles and models have been proposed to interpret their structure and activity relationships (SARs)^3-8^. In this research, the SARs of sweet compounds of widely different chemical families were surveyed from a “trace to the source” view on the molecular organization of their components and their interaction with the sweet taste receptor (STR). This leads to a disclosure of intrinsic connectivity patterns in both sweeteners and STR: charge complementarity and compatibility between components, which afford the complementary sweetener-receptor interaction that induces receptor activation, accounting for the molecular origin of sweet taste. Herein, the analogous topology between glucophores in sweeteners and its counterparts in receptor, and their befitting orientated interaction, which is the common molecular feature of sweeteners, are firstly revealed. This paradigm not only provides a meaningful framework and helpful guidelines for further exploring SARs and molecular modification/design of sweeteners, but also has significant implications to illuminate the underlying mechanisms of molecular origin/evolution of both sweeteners and sweet taste receptors.

## Introduction

Our curiosity for the molecular origin of sweet taste has lasted for a long time, as sensation of sweetness is an undoubted talent for higher animals including humans to respond to their external eating ingredients^1-5^. A basic question still holds: what is the common feature imparting a same taste to sweet compounds with differing types of structures? Seeking could be retrospected to a hundred years ago when multiple hydroxyl groups or a glucophore and an auxogluc were observed as sweet characteristics^2-5^. Since Shallenberger and Acree’s first proposal in 1967 that a sweet agent must include a hydrogen donor (AH) and receptor (B) within a distance 2.5-4 Å^3^, followed by Kier’s supplement that a third hydrophobic site (X) in proper array with AH-B unit is necessary^4^, numerous studies have been performed to elaborate the SARs of various sweeteners ranging from amino acids/sugars to natural (e.g. stevioside) and synthetic compounds (e.g. saccharin) as well as sweet-tasting proteins (STPs)^9-16^. All these results, although mostly sweetener-specific, were derived from the AH-B-X premise (Fig. 1a). Designing new, safe and low-calorie sweeteners is particularly important in medicine to cope with diseases linked to the consumption of carbohydrates^16^.

**Fig. 1.**
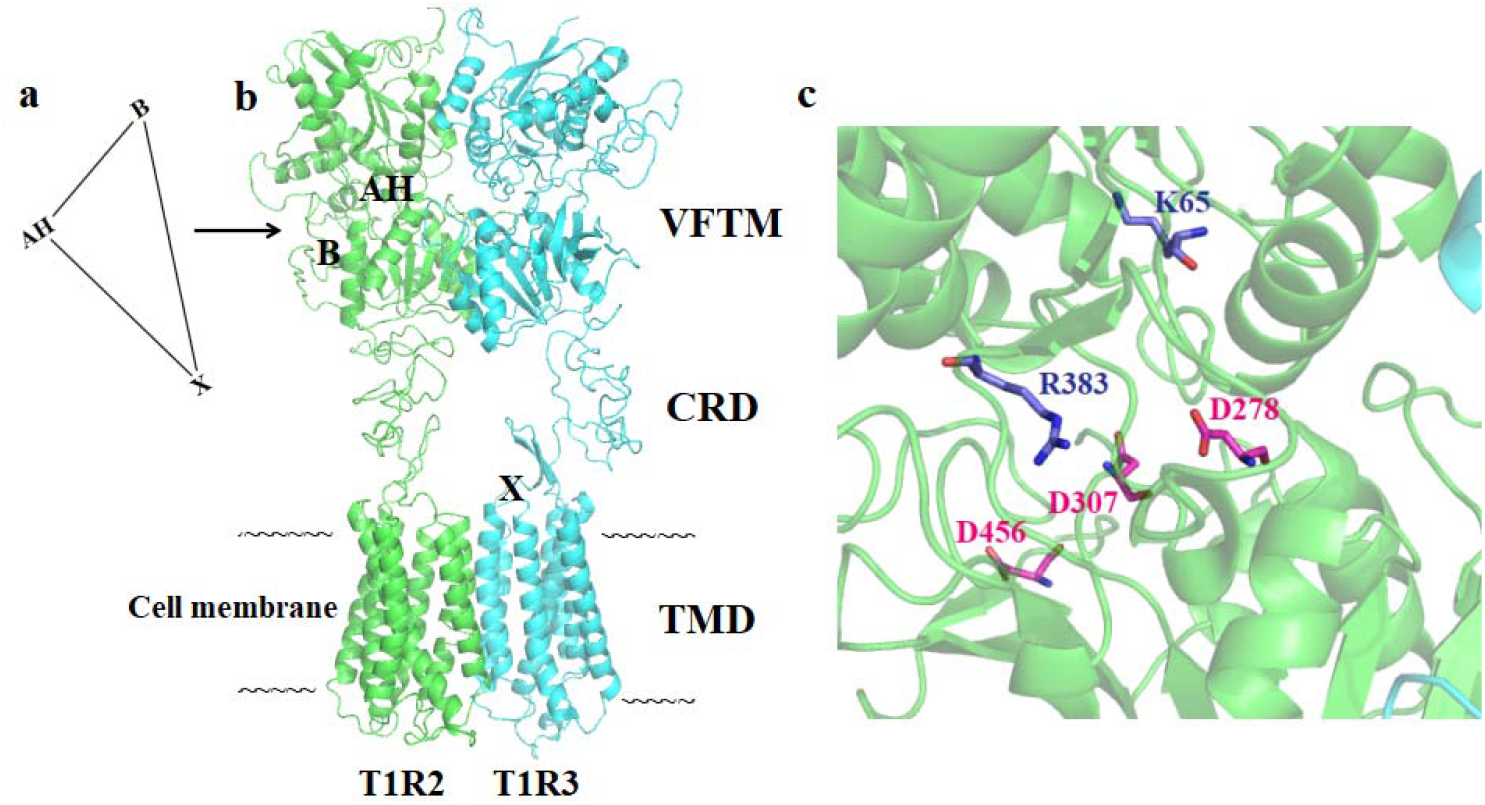
Schematic representation of the glucophores in AH-B-X theory and its counterparts in the sweet taste receptor (STR). **a**, Arrangement of sweetening agents sites in the AH-B-X entity. **b**, The modeled structure of human STR and its potential AH-B-X glucophores interactive counterparts. The T1R2 and T1R3 subunits are colored in green and cyan, respectively, and their VFTM, CRD and TMD are labeled. **c**, Illustration of the proposed “pincer residues” in the orthosteric ligand-binding active site located between the two lobes of T1R2 VFTM, which are shown in stick models and colored in blue for AH counterpart and magenta for B counterpart, respectively. The D456 involved in interaction with STPs is also included. All figures in this article were made with the PyMOL software.

Despite of its wide applicability, however, the AH-B-X paradigm is obviously limited. Some compounds harboring the AH-B-X entity are tasteless and it is hard to interpret the stereoselectivity of chiral amino acids and SARs of high-intensity sweeteners^6,9^. An updated eight-sites attachment theory argued a coexistence of an electrostatic interaction between AH and B with 1-6 additional sites, which are in specific spatial arrangements facilitating the interaction with STR^5^. Furthermore, several models were proposed to elucidate the SARs of representative sweeteners (e.g. aspartame) with crystallology, NMR experiments and theoretical calculations, however, inconsistent conclusions were usually made^11-14^. Moreover, it is difficult to understand the SAR of huge and bulky volume STPs with these models^16^.

Previous studies indicated distinct contributions of three elements (AH, B and X) for the sweet taste, but the results were obviously sweetener-dependent^1,17^. However, it is generally accepted that the charge distribution at the three sites in sweet molecules should be regarded as the primary driving factor for eliciting sweetness. Several investigations correlated the extent of group polarizability and hydrophobicity with the sweetness of different types of sweeteners, but irrelevant results were often obtained^1,7,17^. Although a large number of sweeteners have been investigated with the AH-B-X theory based models, SARs of sweeteners with complicated structures are still unclear, especially for STPs^16^. Evidently, there is no unified theory accounting for the common structural feature of a variety of sweeteners for generating sweetness, that is, the molecular origin of sweet taste.

Since the functional identification of human STR in 2001^18^, substantial research has steered to the interaction details between sweeteners and STR to elucidate their SARs. All these results verified the previous assumption that the diverse sweeteners act on a common receptor^19-20^. In spite of absence of authentic structures of STR and its complexes with sweeteners hitherto, molecular simulations and functional mutagenesis analysis have uncovered a landscape of sophisticated and diversified interplay modes between sweet ligands and involved residues of STR, oversimplifying the AH-B-X paradigm^20^. A uniform mechanism imparting the sweetening power, as exemplified by the close encounter of small molecular weight sweeteners and long-distance communication of large volume STPs with STR, remains elusive.

The structure of heterodimeric human STR (T1R2/T1R3) has been widely acknowledged which includes an extracellular N-terminal Venus flytrap module (VFTM) and a heptahelical transmembrane domain (TMD), which are linked by a cysteine-rich domain (CRD). A relatively tiny intracellular domain (ID) is involved in interaction with the G protein (Fig. 1b)^19^. Receptor activation is initiated by a conformational closure of two lobes of VFTM upon sweet ligand binding, followed by a transmission of induced signal along CRD and TMD to ID. Previous studies have shown that many sweeteners enter the pocket composed of the two lobes of VFTMs, while some bind to TMD of T1R3, stabilizing the activated state^20^. Elucidating the mechanism of ligand-triggered receptor activation is essential for understanding the SARs of sweet compounds as well as their structure-based molecular modifications/designs^2^.

In this report, through generalization and analysis of the SARs of various sweeteners, a novel unified model is proposed to account for the common molecular determinant for sweet taste. It underlines that the intrinsic stereochemical organization of sweeteners, which naturally couples STR activation through counterparts fit, is essential for eliciting sweetness. This mechanistic insight not only provides helpful principles for designing novel sweeteners, but also presents far-reaching implications for decoding the strategies of nature in designing biological perceptions.

## Results

### A unified molecular theory of sweet taste

Inspired by a systematic view of molecular assembly, the SARs of sweeteners were resurveyed in a different way. Specifically, emphasis is on the mechanism that the global property of compounds depends on the intrinsic connectivity pattern of their components^21^. Building on this intuition, topological structures analysis underlines an interrelationship between components in sweet molecules, which is a common molecular feature for generating sweetness. This relationship can be divided into two subcategories.

First, the AH-B entity, the importance of which for eliciting sweetness has been validated in much research, embodies a significant pattern in molecular organization: charge complementarity (considering most radicals in cation or anion form being detrimental). An incongruity between positive and negative charges or their separation over certain range could break this harmony, leading to destruction or decline of sweetness^2-5^. Elimination of the radicals by intramolecular complementation satisfies the harmonious states and stabilizes the advantageous conformation of molecules. This zwitterion and charge/electrostatic complementary characteristic, which is conspicuously differentiated from other compounds in chemical reactions involving bonds breakdown or groups transfer, has been delineated as an essential prerequisite for eliciting sweetness in abundant SAR studies of sweeteners^1-8^.

Second, previous research indicated that proper spatial arrangement between zwitterionic AH-B and X moieties in sweeteners, constrained as their specific conformers, serves as a necessity for sweetness^8,11^. There has also been evidence that besides steric coordination, the length and size of X play key roles in modulation of the intensity of sweetness^1-2,4-8^. Accordingly, another relationship, compatibility between hydrophilic zwitterion and hydrophobic X moiety can be claimed for sweetening performance. This stereochemical pattern, albeit in diversified modalities and sweetener-specific, is not only critical for conformation preference, but also yields stereoselectivity upon receptor binding. This point is vindicated by the hydrophilic/hydrophobic balance in a lot of SAR studies of sweeteners^2,8,17,22-25^.

Next, it is logical to ask whether this feature in sweeteners is relevant to their interactions with STR. The modeled structure of STR consists of two heterogenous subunits, both with a hydrophilic residues rich VFTM and an inner hydrophobic TMD^20^. Strikingly, by analogy with the AH-B-X unit, the overall receptor configuration can be likened as a AH-B-X recipient counterpart, defining a gross orientation and stereoselectivity of sweeteners (Fig. 1b). This formulation is supported by several evidences: (a) The identified “pincer residues” at the upper lobe (basic K65 and R383) and the bottom lobe (acidic D278 and D307) of T1R2 VFTM^26^, which is an orthosteric binding site for majority of sweeteners, can be ascribed to interactive counterparts of AH and B moieties of sweeteners, respectively. A comparable role of T1R2 D456 involved in STPs binding was also described^27^, and there are other potential residues with this behaviour (Fig. 1c)^18-20^. Consistent with this setting, alteration of the orientation of zwitterion completely destroys sweet taste^2,8^; (b) Distinct roles of two subunits involved in hydrophilic (T1R2) and hydrophobic (T1R3) interactions have been demonstrated^12,19,20,27^. It was suggested that the ligand-triggered signal in T1R2 VFTM transmits along the VFTM, CRD and TMD of T1R3, and the latter is a classical allosteric site for most sweet modulators^28^. Accordingly, the T1R3 TMD is akin to a X counterpart, and together with T1R2 VFTM can generally be regarded as a AH-B-X recipient analogue. Evidence for this topological assignment comes from the early argument for a strictly apolar nature for the side of sweet ligands facing the spatial barrier in a narrow cleft shape of active site^11^. This feature is convinced by almost identical orientation of X portions to TMD of comparable AH-B-X entities of ligands in homologous metabotropic glutamate receptors (mGluRs)^12^, an analogous heterodimeric GABA_B_^29^ and docking simulations of sweeteners with functional mutants validations^20^, even for ligand binding at the T1R3 TMD (Supplemental Figs. 1-4); (c) The hydrophilic/hydrophobic partition tendency facilitates the fitness in the interaction between glucophores and their receptor counterparts^12,23,30^. This scheme strongly suggests that only one of enantiomers is capable of interaction with STR since the receptor has chiral character^8,12^, which is in accord with its well-described stereoselectivity for numerous sweet molecules. (d) Most importantly, the topological arrangement of AH-B-X entity in all sweeteners is stereochemically complementary with the counterparts in receptor, as shown in the following validation of this model by evaluating the SARs of different sweeteners.

This intrinsic analogous and complementary relationships between sweeteners and STR with both polar and apolar fit, could be implicated in the molecular origin of sweet taste.

### SAR analysis of sweeteners in different chemical classes

#### Amino acids

It is not completely understood why most D-amino acids taste sweet while L-amino acids not, because the previous virtual “spatial barrier” using Leu is hypothetical, and its applicability for other amino acids is unknown^8^. Moreover, it is unclear why X moiety of sweet D-amino acids must be at least as large as an isobutyl group of Leu. The new model presents a rational explanation. As shown in Fig. 2a-f, if the AH-B moieties of both D- and L-amino acids are identically posed to interact with the B-AH counterparts in receptor, the X moieties of sweet D-isomers of Leu, Phe, His, Tyr and Try adopt suitable orientation toward the receptor X counterpart (Fig. 2i), while the L-isomers do not. This comparison also discloses a reasonable cause for the highest sweetness of Try (≥5-fold than others) due to its superior hydrophobicity of X and orientation to TMD. The accidental sweet L-Ala and nonsweet D-Ala are also interpreted. For amino acids with less side-chain than that of Leu like Val, X moieties in both D- and L-isomers show no appropriate orientation to the hydrophobic portion of receptor. Because of the achiral character of Gly, both its D- and L-isomers are sweet (Fig. 2e-h). It is easy to see that the binding site in T1R2 VFTM can accommodate the sweet molecules with the requisite orientation. Collectively, specific conformer of amino acids with fit coordination to their corresponding counterparts in receptor is thus crucial for eliciting sweetness.

**Fig. 2.**
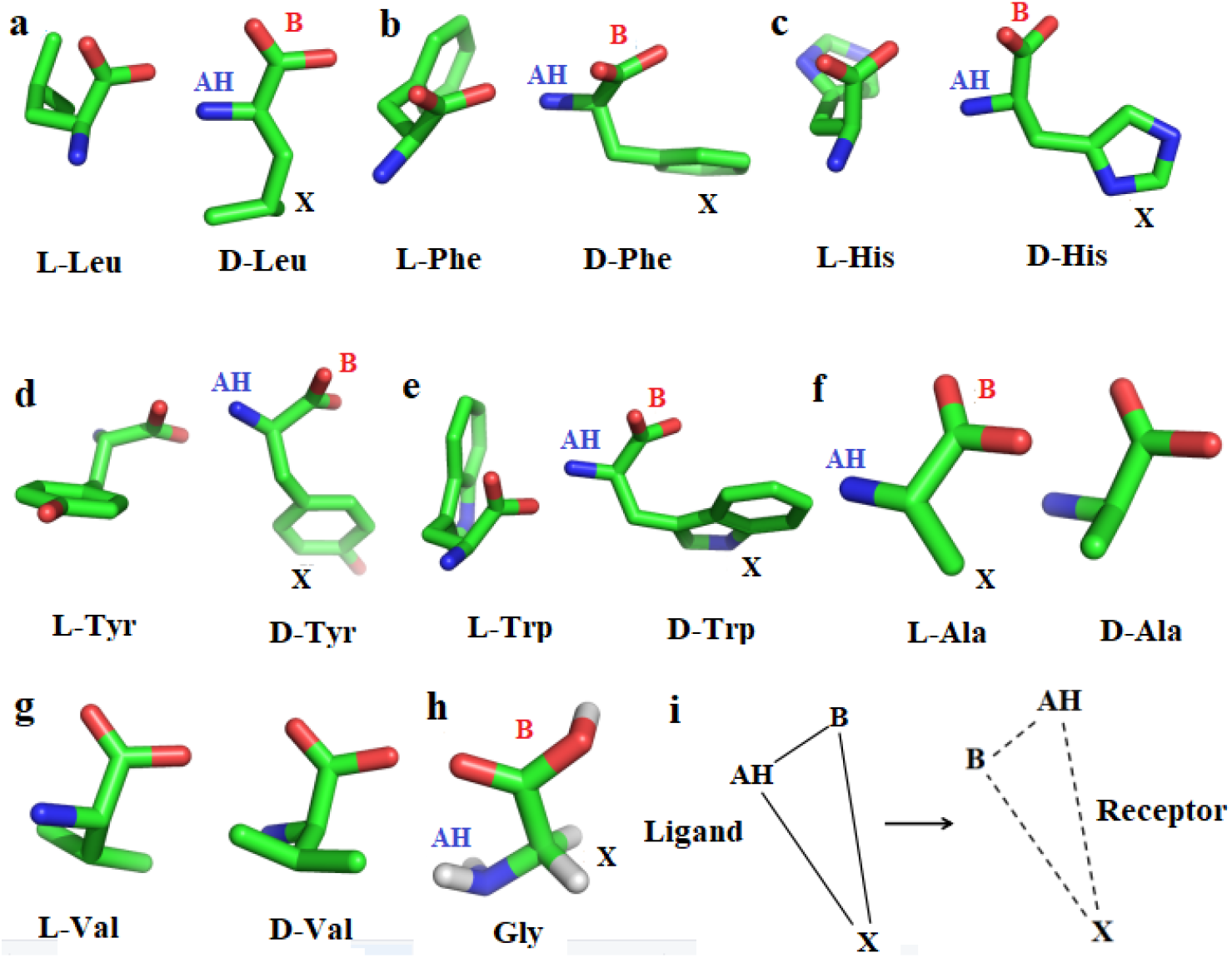
Molecular structure of the enantiomeric taste amino acids. The stereomeric drawing of three dimensional structures of amino acids are shown for visualization. The AH-B-X entity of the sweet taste active isomers are denoted. **a**, Leucine. **b**, Phenylalanine. **c**, Histidine. **d**, Tyrosine. **e**, Tryptophan. **f**, Alanine. **g**, Valine. **h**, Glycine. Hydrogen atoms are added for this amino acid to indicate its X part. The C, N, O and H atoms are shown in green, blue, red and white colors respectively throughout the article. **i**, Illustration of the AH-B-X sites of amino acids and its counterpart of STR.

#### Aspartame

Two distinct bioactive conformations, folded (L shape) or extended form of a well-studied sweet dipeptide aspartame (L-Aspartyl-L-phenylalanine methyl ester) and its analogues were proposed by two independent groups, respectively^6,12^. The chirality selection for sweet taste is consistent to the present model (Fig. 3a). The topological arrangement of glucophores counterparts in STR, functional differentiation of distinct domains between T1R2 and T1R3^12,20,27^ and signal transmission from T1R2 VFTM to T1R3 TMD^28^ strongly argues that the receptor adopts an allosteric trans-activation mechanism that is unique for heterodimeric class C GPCRs^31^. However, conformations of both peptidic ligand and receptor are very flexible, and bioactive conformer of aspartame could be an ensemble with both L shape and extended types^32^. An equilibrium between active L shape and extended conformers could derive from the mutual efficacy in allosteic modulation of receptor activation between T1R2 and T1R3. Alternatively, there could be a conformational switch between two conformers accompanied with receptor activation process, which coincides with the signal transmission direction from VFTM to TMD (Fig. 3b). In support of this, an aspartame analogue with a crown-ether adopted virtually all extended conformations in apolar environment^30^, and ligand reorientation to TMD has been found in other class C GPCRs^29^. Intriguingly, the successful docking of each conformer in active site also reveals a substantial conformational transformation (Fig. 3c,d), and further molecular dynamics analysis could provide more informative details. In both cases, the previously defined relatively broad range of torsion angles requirement for hydrophobic part of aspartame is in accord with the general topological orientation constraints for its X portion toward TMD space. This viewpoint is favored by the fact that four stereoisomers of monatin, which is suggested to interact with receptor in a similar manner as aspartame, all exhibit sweet taste but the (2R,4R)-isomer displays the most intense sweetness^33^.

**Fig. 3.**
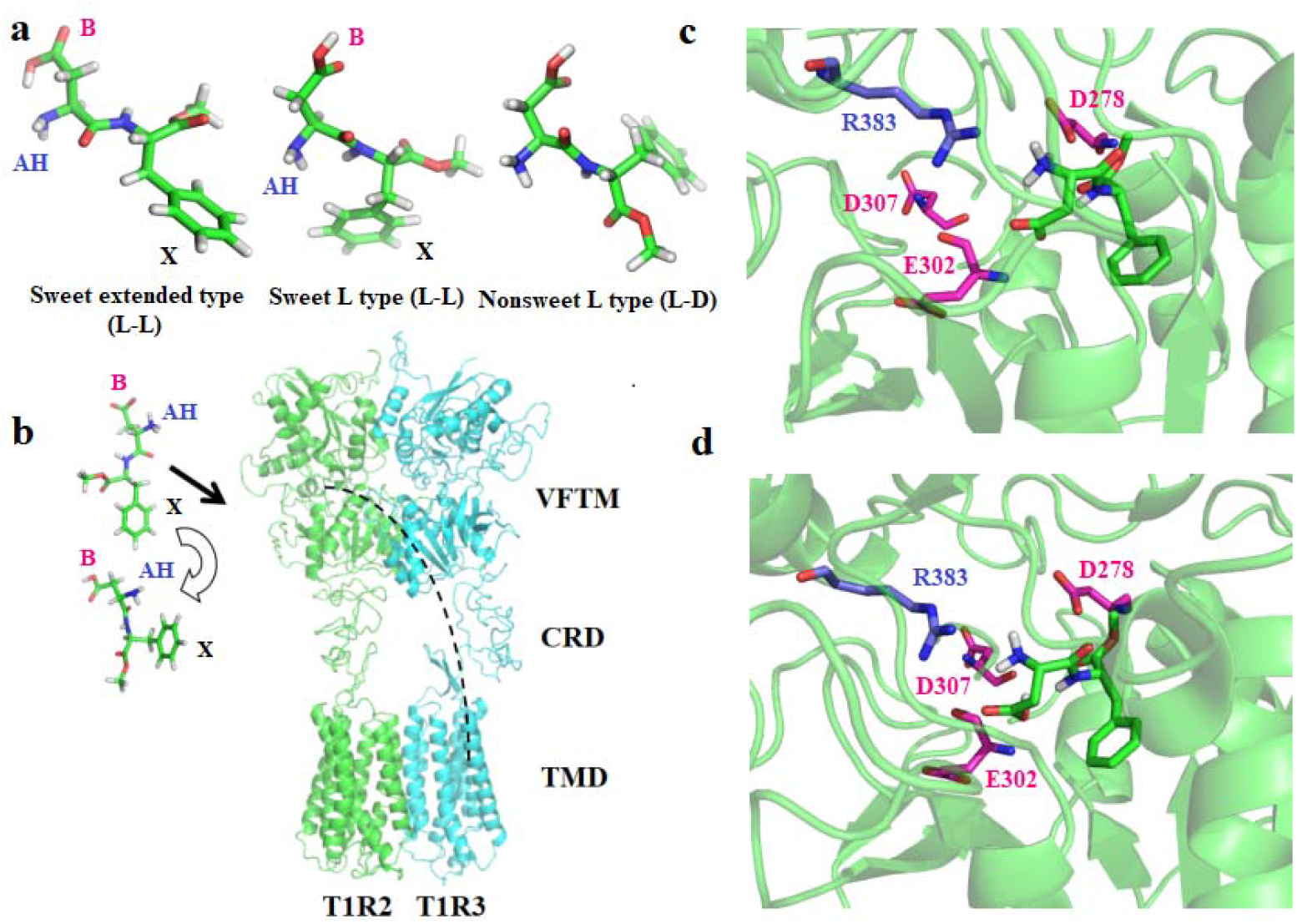
Modeling of the molecular interaction between aspartame and sweet taste receptor. **a**, Three dimensional structure of aspartame (L-aspartyl-L-phenylalanine methyl ester, L-L isomer) with two active conformations (L shape and extended forms) which taste sweet. The nonsweet L-D isomer is added for comparison. **b**, A proposed conformational transformation of aspartame during the activation of STR. The extended and L shape conformers with AH-B-X sites annotation are shown on the left, and a dotted curve indicating the transmission pathway of triggered signal upon ligand binding is shown on the right. **c**, Docking aspartame (L shape form) into the T1R2 VFTM of human STR. The aspartame molecule and involved residues as AH and B glucophores counterparts within the pocket are shown as stick models and their backbones are colored in green, blue and magenta, respectively. **d**, Docking aspartame (extended form) into the T1R2 VFTM of human STR. Same representation is shown as **c** and it is noteworthy that the two conformers in the active site of STR adopt almost identical orientation to T1R3 TMD, probably via a conformational transformation from the extended form to L shape form during the activation of STR.

#### Other typical sweeteners

Previous reports indicated that conformational richness and intramolecular hydrogen bonds in sugars provide their contact points with receptor^34^. This conception can be extended to other plant-derived sweeteners such as perillartine, aniline derivatives, diterpenoid stevioside and neohesperidin dihydrochalcone (NHDHC). Because the substitutes appear significant influences on sweet properties of these molecules^7,17^, RESP (restrained electrostatic potential) charge analysis was performed to unveil their electrostatic potential distribution. The results afford a reasonable depiction of their candidate glucophores. It is clear that the chiral D-Fructose shows same sweetness as its L-isomer because the C_3-5_ configuration has no affect on the AH, B and X sites assignment as OH_1_, OH_2_ and C_6_, respectively^34^ (Fig. 4a). Besides, electronic structures of perillartine and 5-Nitro-2-propoxyaniline (Ultrasweet) reveal alternative AH, B and X glucophores, as previously suggested^17^ (Fig. 4b, c). For stevioside, because the aglycon has been proved to be unnecessary for sweetness elicitation^35^, two units are found to be AH-B-X candidates, and the importance of C_16_-C_17_ methylene double bond for sweet taste in previous synthetic chemistry ascertainment is also clarified^9^ (Fig. 4d). The results of NHDHC is also consistent with the former allocations of its AB-H-X units^24,37^, however, similar to stevioside^36^, other candidate glucophores, which may be engaged in the enhancement of total sweetness, cannot be excluded (Fig. 4e). Although different sweeteners display obviously varied chemical constitution and spatial configuration, all these compounds exhibit considerable conformance with the topological orientation requirement in the present counterparts fit model.

**Fig. 4.**
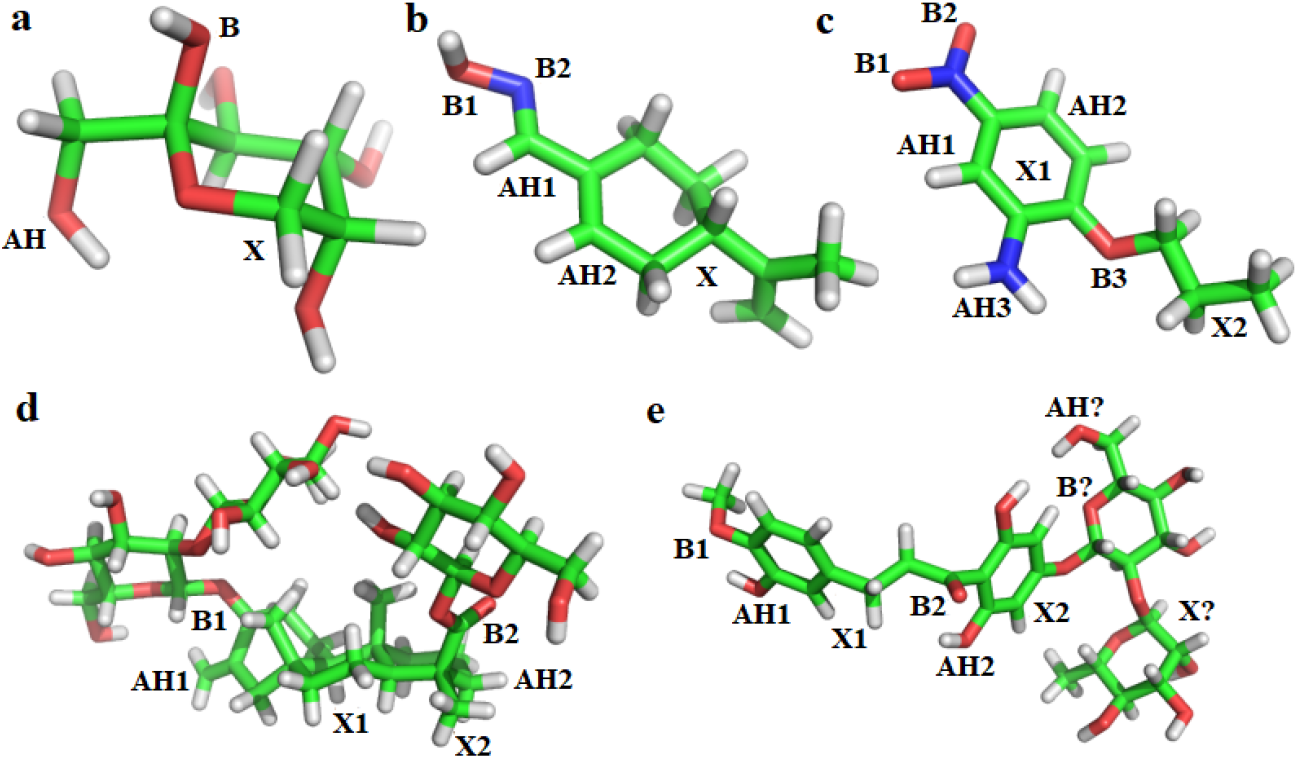
The AH, B and X glucophores assignment of representative sweet compounds. The glucophores candidates are deduced based on the electrostatic potential distribution calculated with the RESP (Restrained ElectroStatic Potential) approach. **a**, L-Fructose. **b**, Perillartine (1-perillaldehyde α-antioxime). Potential multiple AH-B-X glucophores are numbered. **c**, 5-Nitro-2-propoxyaniline. **d**, Stevioside. **e**, NHDHC (neohesperidin dihydrochalcone). The hypothetical AH-B-X unit candidate in the β-D-sophorosyl aglycon (right) is denoted.

It is evident that there could be multiple potential AH, B and X sites in these sweeteners, and their conformational diversity could confer multifarious interplay modes with receptor. In line with this finding, previous investigations indicated that number of polyhydroxy and hydrophobic sites within a limited range are positively correlated with the sweetening potency^1,22^. Furthermore, extra X sites are shown to be involved in the modulation of sweetness^5,17^, and alternative AH-B-X sites with a more optimum hydrophile/lipophile ratio are suggested as structural signatures of sweeteners^17,37^. Therefore, it seems that combination of distinct (AH-B-X)s could enhance sweetness in concert, provided with their appropriate spatial arrays. A synergetic effect of glucopheres with multivariable nonlinear relationships can thus be envisaged. However, predicting the performance of this ensemble could be considerably complicated due to the induced and delocalized effects in the whole molecule, their topological disposition and tremendous conformational flexibility. Nevertheless, the intrinsic connectivity pattern in AH-B-X units and their complementary fit with the counterparts in receptor, mediated by intramolecular organization and intermolecular interaction respectively, play a crucial role for arising of sweetness.

#### Sweet-tasting proteins

It is still mysterious that the large-sized STPs interact with and activate human STR. Employing the present model, a remarkable analogous pattern is also revealed: the natural monellin harbors two chains A and B, and residues located at the N and C terminals of chain B are mostly flexible and charged^16,27^. Mutations of these sites modify a zwitterion-like potency generally result in obvious alteration of sweetness (Supplemental Tables 1 and 2). On the other hand, the C terminal of chain A is enriched with hydrophobic residues, dominated by a distant poly-(L-proline) II (PPII) helix, whose deletion is detrimental for sweetness^38^. This architecture renders an AH-B-X resemblance of its glucophores (Fig. 5a). Moreover, its overall wedge shape is comparable to the AH-B-X configuration in nonproteic sweeteners^27,39^. Such pattern is also present in brazzein and thaumatin (Fig. 5b, c). Functional mutants results of surface residues can be relied to map the potential glucophores for all three STPs. For instance, a number of basic residues could be ascribed to the AH glucophores of the three STPs, while a few acidic residues could be appointed as B glucophores, in agreement with the statement that the surface of STPs is generally positive charged^27^. Hydrophobic glucophores PPII helix region of monellin, Y39 and disulfide bond C_16_-C_37_ of brazzein and F80 of thaumatin could also be proposed, and there could be other potential glucophores in these STPs, indicating a complex map of glucophores distribution (Supplemental Table 2). Nevertheless, it is indicated that adjustments of either zwitterionic or hydrophobic portions of three STPs lead to potentiation or detriment of the overall efficacy of glucophores with increased or decreased sweetness, suggesting a similar interactive fashion of STPs with the counterparts in receptor as that of nonproteic sweeteners.

**Fig. 5.**
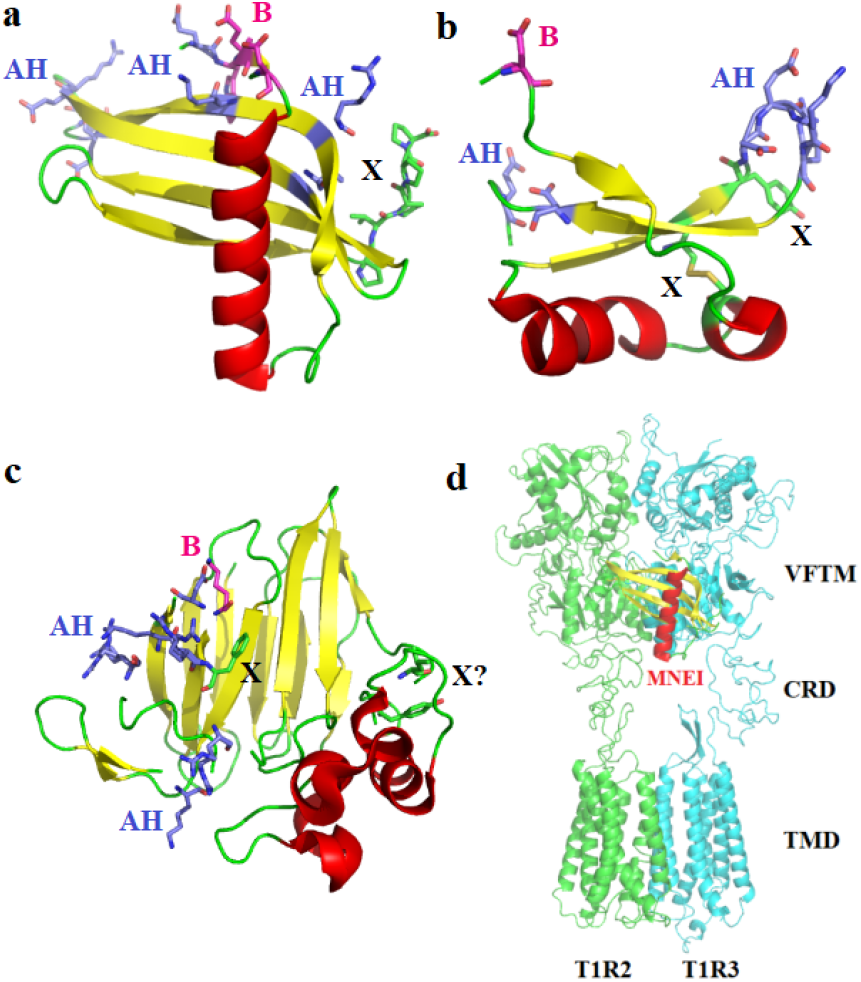
The graphic illustration of the structures of sweet-tasting proteins and their potential glucophores. The α-helixes, β-sheets and loops in the cartoon model structures are colored in red, yellow, and green, respectively. The glucophores are assigned based on the functional site-directed mutagenesis results. **a**, Monellin. The structure of a single-chain monellin (MNEI) (PDB ID: 1FA3) is adopted due to its same spatial structure as that of native monellin, and its candidate glucophores residues are shown as follows. AH: E2/R39/K43/E50/R53/E54/Y65; B: W3/E4/D7; X: hydrophobic poly-(L-Proline)II helix region (P92-P96). The W3 and E4 are assigned as B portion because of their adjacency with the flexible and charged N-terminal and their substantial decrease of sweetness upon mutation or vital role in protein expression/folding, respectively. The main chain O atom of W3 could interact with the AH counterpart residues in receptor. **b**, Brazzein (PDB: 4HE7). AH: D40/E41/K42/R43/D50/E53; B: D2; X: C16/C37 and Y39. **c**, Thaumatin (PDB: 3AL7). AH: D21/K67/R76/K78/R79/R82; B: K19; X: F80 and Y183/L185 (hypothetical). D21 is assigned as an AH glucophore site becuase its potential interaction with the acidic residues in receptor, while K19 is assigned as a B glucophore site because it is hydrogen bonded with D21 in both crystal structures of wild-type thaumatin and its D21N mutant. F80 is assigned as X glucophore owing to its potential role in epitope study. The hypothetical X glucophores Y183/L185 without experimental evidence are denoted. **d**, Interaction model between the MNEI and human STR.

This scheme is reinforced by the long-distance electrostatic interaction between surface residues of STPs and VFTMs of receptor^16^. Similar to stevioside and NHDHC, there are multiple potential AH, B and X sites in STPs, which are in accord with the so-called multi-point interactions with STR and their intensively sweet taste^39^. However, these charged residues unevenly spread across the protein surface, and their non-uniform distribution has been demonstrated^40^. The assignments of glucophores aid a coarse representation of interaction (Fig. 5d), in which the binding region of MNEI in STR is alike to that in previous fuzzy representations of STP-STR complexes but with different orientation, most likely due to the low resolution modeling mode, as previously claimed^27^. This interaction mode is also consistent with the finding that crucial electrostatic interactions take place between STP and T1R2, whereas most hydrophobic interactions belong to the T1R3^27^, and the conclusion that the CRD of T1R3 is essential for the sweetness of STPs^41^. The adjacency of two hydrophobic portions renders a rational explanation for the extraordinary sweetening potency of STPs than other sweeteners (Fig. 5d). Thus, the counterpart fit model for small-size sweeteners is also applicable for STPs, although they have different binding modes. However, the abundant residues and extreme conformational flexibility of STPs raise a great challenge to precisely clarify their SARs details, especially one considers the condition in studying dipeptide sweetener aspartame and its analogues^6,12^.

Taken together, it can be proposed that a common pattern of intrinsic intramolecular organization with both charge complementarity and components compatibility in sweeteners, which is essential for their spatial orientation interaction with the counterparts in receptor, accounts for the molecular origin of sweet taste. However, the diversified molecular compositions and conformations of both sweeteners and receptor as well as their topological constraints make their interactions be very sweetener-specific, giving rise to a variety of sweet properties of sweeteners with different chemical structures.

### Interaction between sweeteners and STR

How do the sweeteners interact with the STR structurally, especially one considers the bulky STPs? The popularly accepted wedge model proposes that proteins bind to an external site of STR via their surface electrostatic complementation^16,27^. All SAR models underline a necessity to recognize the AH-B-X triad as a unitary agonist to probe its overall performance in interaction with STR. Upon sweet compounds binding with receptor, the interactive hydrogen bonds, hydrophobic or Van der Waals’ forces could spontaneously induce charge redistribution and orientation readjustment in both molecules, and their linkage lead to an advantageous conformation complex, stabilizing the active form of receptor^12,20,27^. The AH-B moiety of small-sized sweeteners can easily form charge complementation (or hydrogen bond) with its receptor counterpart residues, while the X moiety may bind with a hydrophobic partner but also probably contact with receptor via dispersion interaction^4,5,7^. This interplay mode can be extended for STPs. The long-distance interaction between proteins has been well identified^42^. Propagation of conformational changes over long distances in STPs, mediated by intramolecular allosteric communication networks, has been experimentally verified^38^. In this way, intrinsic disorders in both proteins could optimize their allosteric couplings^43,44^. Furthermore, mutants of numerous surface residues of STPs indicate a rather flexible binding interface^27,39^. Therefore, the glucophores in STPs are apt to exert their affect on receptor via long range interactions. In this regard, the distances between AH, B and X proposed by Shallenberger and Kier should be too strict^3-5^. No matter what kinds of interaction forces operate, the complementary fit between sweeteners and receptor, which derives from their intrinsic molecular organization, is paramount for elicitation of sweetness.

It can also be suggested that receptor activation by stevioside and NHDHC acts in a similar manner and at an intermediate level between simple amino acids/sugars and STPs, as manifested by their moderate sweetness compared to the other two classes of sweeteners. However, an analogues interplay pattern with receptor can be derived for all sweeteners.

## Discussion

This study reveals a common molecular feature for generation of sweetness, which can be regarded as a unified molecular theory of sweet taste. The reliability of this model is validated by its significant heuristic power, as shown in the interpretation of SARs of various sweeteners. It is not contrary to former theories. Moreover, it conforms to some previous models but raises deep insights in a “trace to the source” and inclusive manner^21^. The results highlight that stereochemical complementarity and compatibility between glucophores is the intrinsic common attribute of sweeteners.

Additionally, it also promotes our understanding of the SAR of STR as well as its molecular origin. The receptor belongs to class C GPCRs, most of which are homodimers. However, the overall architecture of heterodimeric T1R2/T1R3 displays asymmetry. This configuration could suitably accommodate sweeteners with widely different chemical structures. Herein, the firstly reported analogous topology between glucophores and its counterparts in receptor as well as their complementary fit in interaction suggest an inherent correlation in probable congenerous origins of sweet compounds and STRs, which could also be associated with the mechanism of molecular evolution of STRs in species^45^.

This report highlights a necessity to view the sweet molecule as a system, with each glucophore being distinct elements. Other elements (e.g. atoms or bonds), which are involved in regulating the properties of glucophores, can also be included. All elements are correlated by high-order interactions to constitute a network responsible for the elicitation and performance of sweet taste. Therefore, research to dissect the intricate connectivity relationships of these elements would be informative for further probing the special model for specific type of sweeteners. To achieve this aim, sweetness must be regarded as an additive property which derives from interlinked and synergistic actions of different components with all molecular features of each component included as a whole^21,46^. Therefore, principles in system biology and network science could be transplanted to this promising research area^47^. It also hints that most planar features of networks should be reevaluated under spatial topology circumstances. This scheme could also be applicable in further research on other sorts of tastes (e.g. umami)^48^.

In conclusion, this work raises a unified model to illuminate the mechanism of molecular origin of sweet taste, which provides a novel framework and meaningful strategies for further SAR investigation and molecular design of sweeteners. This seminal paper, albeit in a general way, also presents significant implications for deeper understanding how nature design chemical reactions to elicit different categories of taste, and even other physiological and neural perceptions.

## Methods

### Molecular modeling

Based on the activation mechanism of class C GPCRs mGluRs and GABA_B_, the ligand-bound conformation of human STR should be an “Acc” (active closed–closed) or “Aco” (active closed–open) form^12,27^. This allows to select the active form of GABA_B_ as the template (PDB: 6UO8) to model the “Aco” (active closed–open) form of full-length STR structure, because the heterogeneous nature of two subunits of this template could well reflect the distinct attributes between T1R2 and T1R3 of STR^29^. A multiple sequence alignment of human STR and GB1 and GB2 was generated by ClustalW^49^, and human T1R2 (Genbank accession no: Q8TE23) and T1R3 (Q7RTX0) display around 25%∼27% overall sequence identities with GABA_B_ GB1 and GB2. A hetero target model was built using the online SWISS-MODEL program (http://swissmodel.expasy.org/). The quality of model was evaluated by the QMEAN scoring function^50^ and Verify3D program^51^ with acceptable scores.

### Quantum chemical calculation of RESP charges

RESP (restrained electrostatic potential) charge analysis of sweet compounds was carried out with the Gaussian 09W program^52^ and AmberTools 20 program^53^. The three dimensional structures of compounds were downloaded from the NCBI PubChem Compound database (https://www.ncbi.nlm.nih.gov/), except those of D-fructose and stevioside were extracted from the CCDC (Cambridge Crystallographic Data Centre) database (https://www.ccdc.cam.ac.uk/). The extended conformer of aspartame was drawn with the GaussView 6.0 software. All structures were optimized at the HF/6-31G(d) level. The ESP (electrostatic potential) calculations were performed by the DFT (density functional theory) method at the B3LYP/6-311G(d, p) level, and RESP charges were then obtained using antechamber in the AmberTools.

### Docking simulations

An automatic molecular docking program in Yinfo Cloud Computing Platform (https://cloud.yinfotek.com/) was applied to construct the the complex of aspartame molecule (extended and L type conformers respectively) and the modeled human STR, with the previously well-characterized aspartame-receptor interactive residues (S40, Y103, D142, S144, S165, S168, Y215, P277, D278, L279, E302, S303, D307, R383 and V384) as the constraints^54-55^. The Autodock Vina protocol and the flexible docking model were used. The final docked aspartame configurations were selected based on docked binding energies and cluster analysis. Docking of cyclamate into the human T1R3 TMD was performed with the same procedure according to its previously reported binding site with identified residues (Q636, Q637, S640, H641, L644, H721, R723, F730, F778 and L782) as the constraints^56^.

### Recombinant protein expression, purification and mutagenesis

The recombinant vectors pET15b and pET-SUMO harboring the single-chain monellin (MNEI) and des-pGlu1-brazzein genes respectively were constructed as previously described^57,58^. Expression and purification of wild-type and mutants of MNEI and brazzein were performed according to the corresponding procedures. The His-tag sequence of MNEI was then removed with the thrombin to eliminate its probable effect on the protein properties, and the SUMO-tag sequence of brazzein was removed to obtain the taste proteins. All variants were constructed according to the standard PCR-based mutagenesis strategy (Stratagene) and verified by DNA sequencing.

### Sensory evaluation

Evaluations of the sweetness thresholds of recombinant wild-type STPs and their mutants were performed with a double-blind taste assay, as previously described^58^. Relative sweetness were calculated as 100% of the wild-type threshold/mutant threshold. Comparison of the sweetness of wild-type STPs and their mutants^59-70^ is detailed in Supplemental Table 2.

## Data availability

Source data are provided with this paper. The datasets generated during and/or analysed during the current study are also available from the corresponding author on reasonable request.

## Acknowledgements

This work was supported by the National Natural Science Foundation of China (31970935).

