## Supplemental Files for "A unified molecular theory of sweet taste: revisit, update and beyond"

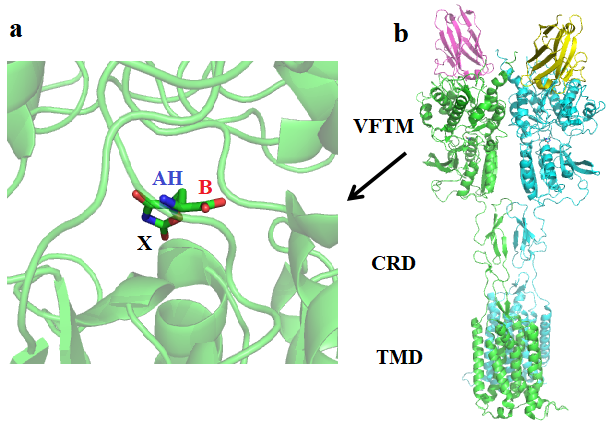

**Supplemental** **Fig. 1. Complex structure of the human metabotropic glutamate receptor 5 (mGluR5, class C GPCR) with the ligand L-quisqualate (PDB:** **6N51).** **a,** The stick model of L-quisqualate and topological orientation of its AH, B and X glucophores analogues. **b,** A cartoon representation of the full-length mGluR5 structure. The two subunits of mGluR5 (chains D and A) are colored in green and cyan and two chains (B and C) of nanobody 43 are colored in magenta and yellow, respectively. It is shown that the X portion analogue of ligand in chain D orientates to the TMD of chain A of the receptor.

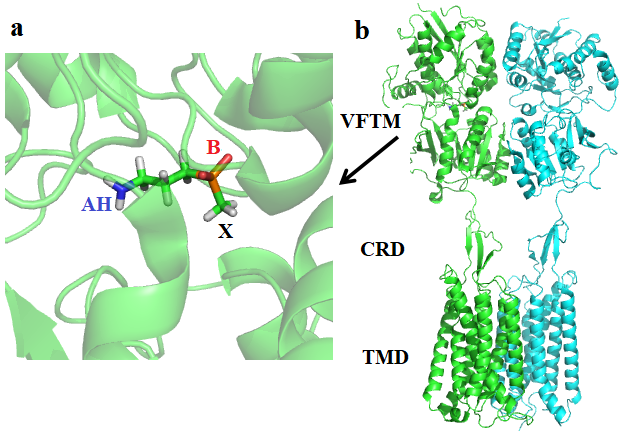

**Supplemental Fig. 2. Complex structure of the human Metabotropic γ-aminobutyric acid receptor (GABA_B_, class C GPCR) with the ligand 3-aminopropyl(methyl)phosphinic acid (SKF97541, PDB: 6UO8).** **a,** The stick model of SKF97541 and topological orientation of its AH, B and X glucophores analogues. Hydrogen atoms are added for visualization. **b,** A cartoon representation of the full-length GABA_B_ structure^29^. The heterodimer is composed of two subunits GB1 and GB2 which are colored in green and cyan respectively. Similar orientation of the X portion analogue of ligand to TMD is shown as those in mGluR5 and STR.

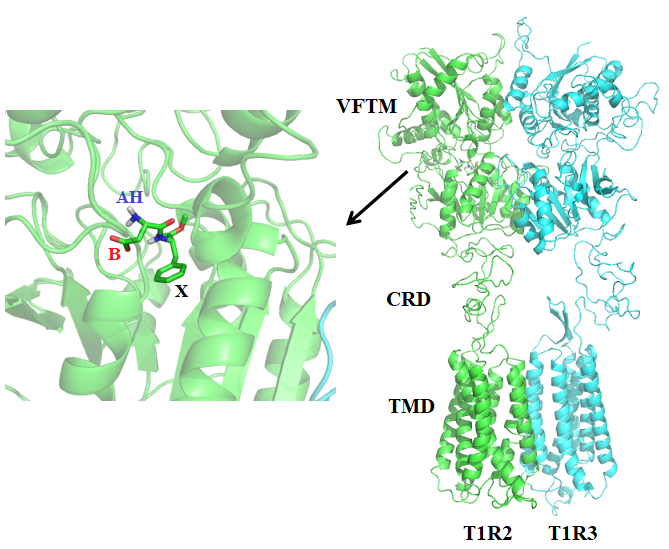

**Supplemental Fig. 3. The complex model of human T1R2/T1R3 and a well-studied synthetic sweetener aspartame. a,** The stick model of aspartame (L shape conformer) and topological orientation of its AH, B and X glucophores based on the docking simulations. **b,** A cartoon representation of the modeled full-length human T1R2/T1R3 structure. The orientation of the X portion (the C-terminal benzyl side chain of phenylalanine and methyl ester) of aspartame to T1R3 TMD is shown as those in mGluR5 and GABA_B_. Although the structures of ligands and receptors are different in the three class C GPCRs, their X moieties (or analogues) of ligands in VFTM of one subunit are identically orientated to the TMD of another subunit in a general way^29,54,55^.

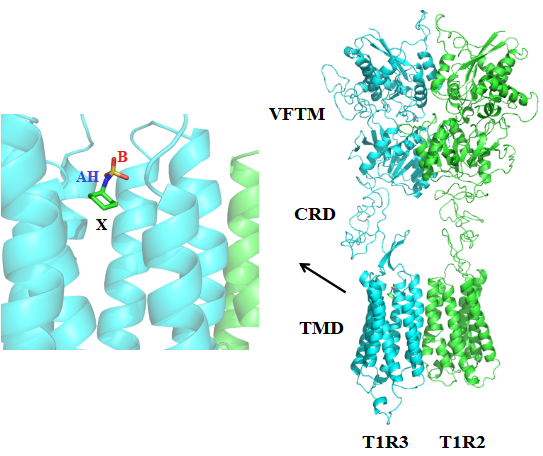

**Supplemental Fig. 4. The complex model of human T1R2/T1R3 and a synthetic sweetener** **cyclamate binding at T1R3 TMD. a,** The stick model of cyclamate and topological orientation of its AH, B and X glucophores based on the docking simulations. **b,** A cartoon representation of the modeled full-length human T1R2/T1R3 structure. The orientation of the X portion of cyclamate in T1R3 TMD is shown, indicating that its hydrophobic X moiety adopts identical topological arrangement as those of aspartame and ligands in mGluR5 and GABA_B_^29,54-56^.

**Supplemental Table 1** **Sweet taste thresholds the wild-type and variants of sweet-tasting proteins monellin and brazzein.**

| Sweet-tasting proteins | Variants | sweet threshold (mM)^a^ |
| --- | --- | --- |
| MNEI^b^ | Wild-type | 1.1±0.09 |
|  | W3A | 75±6.6 |
|  | E4N | No recombinant protein obtained |
|  | R39A | 20±1.8 |
| des-pE1M-brazzein^c^ | Wild-type | 1.5±0.1 |
|  | D2K | 2.4±0.2 |
|  | D2G | 2.8±0.33 |
|  | D50K | 0.3±0.03 |
|  | E53R | 0.5±0.07 |

^a^ Sweet taste thresholds the wild-type and variants of STPs were evaluated with a double-blind taste assay, as described in Methods. Standard deviation (SD) was calculated from the results of three paralleled experiments. ^b^ The single-chain monellin (MNEI) with the two natural chains being joined via a Gly-Phe dipeptide linker, which shows identical spatial structure and sweet properties as the native monellin. ^c^ A form that lacks the N-terminal pyro-glutamate (pGlu) residue.

**Supplemental Table 2 Potential glucophores of sweet-tasting proteins monellin, brazzein and thaumatin according to the functional variants studies**

| **Sweet-tasting proteins** | **Variants** | **Sweetness (%)^a^** | **Glucophore site** | **References** |
| --- | --- | --- | --- | --- |
| MNEI^b^ | Wild-type | 100 | AH-B-X |  |
|  | E2N | 333.3 | AH | 59 |
|  | E2K | 355 | AH | This study |
|  | W3A | 1.5 | B | This study |
|  | E4N | No protein obtained | B | This study |
|  | D7N | 0 | B | 60 |
|  | R39D or E | 0 | AH | 60 |
|  | R39A | 5.5 | AH | This study |
|  | K43E | 8 | AH | 61 |
|  | E50N | 275 | AH | 59 |
|  | E50K | 297 | AH | 59 |
|  | R53A | 37.9 | AH | 59 |
|  | E54N | 150.7 | AH | 59 |
|  | Y65R | ~165 | AH | 40 |
|  | P96 (deletion) | 15-20 | X | 61 |
|  | P93-P95 (deletion) | 10-15 | X | 61 |
|  | P92-P96 (deletion) | 5-7.5 | X | 61 |
| des-pGlu1-brazzein^c^ | Wild-type | 100 | AH-B-X |  |
|  | D2E | ~133 | B | 62 |
|  | D2K | 62.5 | B | This study |
|  | D2G | 53.6 | B | This study |
|  | D40K | ~200 | AH | 62 |
|  | E41Q | ~150 | AH | 62 |
|  | K42A | ~40 | AH | 62 |
|  | R43N | 0 | AH | 62 |
|  | D50K | 500 | AH | This study |
|  | D50A | 20.7 | AH | 63 |
|  | E53R | 300 | AH | This study |
|  | E53D | 45.2 | AH | 64 |
|  | C16A/C37A | ~26 | X | 65 |
|  | Y39A | ~48 | X | 65 |
| Thaumatin^d^ | Wild-type | 100 | AH-B-X |  |
|  | K19A | 64.3 | B | 66, 69 |
|  | D21N | 160 | B | 67 |
|  | K67A | 5.2 | AH | 66, 68 |
|  | K67E | 3 | AH | 68 |
|  | R76A | 47.4 | AH | 66 |
|  | K78A | 20.2 | AH | 69 |
|  | R79A | 26.5 | AH | 66 |
|  | R82A | 4 | AH | 66, 68 |
|  | R82Q | 4.5 | AH | 68 |
|  | R82E | 0.5 | AH | 68 |
|  | F80 | Epitope study | X | 70 |

^a^ For assessment of the affect of mutations on sweetness, sweetness values are normalized and expressed as the ratio (%) of sweet threshold of wild-type, which is of 100% level, to those of variants. This values give the relative sweetness of variants compared with the wild-type proteins. ^b^ Same as Supplemental Table 1. ^c^ Same as Supplemental Table 1. ^d^ A major form thaumatin I.
